# Integrative Single Cell Multiomic Profiling Analysis Reveals HOX-PBX Gene Regulatory Network Contributing to the Survival of mTOR Hyperactive Cells

**DOI:** 10.1101/2022.09.27.509560

**Authors:** Tasnim F. Olatoke, Andrew Wagner, Aristotelis A. Astrinidis, Minzhe Guo, Erik Y. Zhang, Alan G. Zhang, Ushodaya Mattam, Kyla Chilton, Elizabeth J. Kopras, Nishant Gupta, Eric P. Smith, Magdalena Karbowniczek, Maciej M. Markiewkski, Kathryn Wikenheiser-Brokamp, Jeffrey A. Whitsett, Francis X. McCormack, Yan Xu, Jane J. Yu

## Abstract

Lymphangioleiomyomatosis (LAM) is a rare, debilitating lung disease that predominantly affects women of reproductive age. LAM is characterized by the infiltration of the lungs by abnormally proliferating smooth muscle-like cells of unknown origin via an estrogen-dependent metastatic mechanism. LAM cells carry deleterious mutations of tuberous sclerosis complex (TSC1/TSC2) genes, resulting in hyperactivation of the mechanistic target of rapamycin complex 1 (mTORC1) and ultimately dysregulated cell growth. Sirolimus, an FDA approved mTORC1 inhibitor and current best-choice medication for LAM stabilizes lung function in most LAM patients. However, it requires sustained application and remains inefficacious in some patients. The greatest barriers to finding a cure for LAM include its undetermined origin and unclear underlying pathogenesis. Our study aims to advance knowledge on the origin of LAM, and ultimately serve as a premise for the development of novel therapeutic targets for LAM. Single cell RNA sequencing (scRNA-seq) is a powerful tool in biomedical research that informs gene expression differences at the cellular level and may provide insights into the most fundamental origin of LAM cells. Our scRNA-seq analysis of LAM cells revealed a unique population of cells (LAM^CORE^), expressing uterine-similar homeobox transcription factors (HOX) and Pre-B-cell leukemia homeobox 1 (PBX1), which are absent in normal lung, suggesting that the uterus is the primary origin of LAM. PBX1 is a transcription factor critical for female reproductive tract development and maintenance, and its overexpression is implicated in some female reproductive cancers. In this study we hypothesize that PBX1 promotes survival and lung colonization of LAM (TSC2-null) cells. Using LAM patient-derived cells, we validated the transcriptional profile, gene expression and protein levels of PBX1. We have the first functional evidence that PBX1 and its downstream targets are upregulated in LAM cells. In a mouse model of LAM, we monitored the effect of suppression of PBX1 by short hairpin RNA-mediated gene silencing on lung colonization and tumor growth. We also found that pharmacological suppression of PBX1 attenuates LAM lung colonization and promotes death of LAM cells in vivo and vitro. Our data collectively suggests that PBX1 is a critical regulator of LAM progression.

## INTRODUCTION

Lymphangioleiomyomatosis (LAM) is a rare metastasizing lung disease that affects almost exclusively premenopausal women, and the lung manifestations are characterized by proliferation of abnormal smooth muscle-like cells, leading to the formation of emphysema-like lung destruction. LAM is cause by deleterious mutations in *TSC1* or *TSC2* ^1, 2^, which inhibits the mechanistic target of rapamycin complex 1 (mTORC1). Directly resulting from this discovery, mTOR inhibitors (mTORi), have been shown to stabilize lung function in most LAM patients. However, challenges remain to optimize the treatment of LAM patients. mTORi are only cytostatic, requiring sustained application and have side effects that are exacerbated by prolonged exposure. Thoroughly understanding the features of LAM model would not only improve treatment of LAM but also have major implications for other common diseases with shared characteristics.

LAM lesions are metastatic with uncertain origin. The most definite evidence is that cells within recurrent LAM lesions that arise in the allografts of transplanted LAM patients originate from the recipient, supporting a metastatic mechanism for the disease ^3-5^. Circumstantial data have pointed to a uterine source in part because of the predominance in females, and case series that have described LAM lesions in the uterus of patients ^6-8^, although definitive proof is lacking. The recent application of single-cell transcriptomics in LAM begun to shed light on the cell of origin, the diversity of cell subtypes and functional states within LAM lesions, and the altered gene pathways and transcriptional programs related to disease pathogenesis {Guo, 2020 #8;Obraztsova, 2020 #9;Tang, 2021 #16}. Our analysis of LAM lung scRNA-seq identified a unique population of LAM^CORE^ cells, expressing uterine-specific HOX genes that are not detected in normal lung ^9^.

Female reproductive tract development and diseases are often found to be hormonally dependent ^10-14^. The lack of detailed mechanistic understanding of the roles of female hormones in the LAM pathogenesis has hindered the development and optimization of new therapies for LAM patients. Our single cell RNA-sequencing (scRNA-seq) analysis of human LAM lung discovered that LAM cells express a unique panel of signature genes selectively expressed in normal uterus and known to be important in sex steroid action; these include homeobox transcription factors *HOXA10, HOXA11, HOXD11, PITX2* and Pre-B-cell leukemia homeobox 1 (PBX1) ^9^.

PBX1 is a cofactor of HOX, forming dimers with HOX proteins and determining target gene specificity and the mode of transcriptional regulation ^12^. Particularly relevant to LAM, the HOX/PBX1 dimer has been reported in oncogenesis of other steroid-responsive cancers and has become a potential therapeutic target ^13, 15-17^. One of commonly used reagents to block HOX/PBX1 dimerization is HXR9, a cell-permeable synthetic peptide, that induces apoptosis in cancer models ^16^. The roles of HOX/PBX1 in LAM lesions and the intricate interplay between HOXs and estrogen responses in LAM is completely unexplored. In the present study, we applied single cell multiomic data integration and gene regulatory network approaches in conjunction with functional assays using *in vivo and in vitro* preclinical LAM models to reveal the molecular interplay among HOXs and PBX1 and the functional impact of HOX/PBX1 on the survival of TSC2-null patient-derived cells.

## RESULTS

### Integrative single-cell analysis of gene expression and DNA accessibility identified induction of uterine specific homeobox family of transcription factors in the LAM lung

We previously performed scRNA-seq analyses of lung samples from patients with LAM ^18^ and identified a unique population of cells, termed LAM^CORE^, that were readily distinguished from endogenous lung cell types. LAM^CORE^ cells shared closest transcriptomic similarity to uterine myocytes in both normal and LAM uteri ^18^. In the present study, we generated paired measurements of single cell chromatin accessibility and gene expression using 10x multiome data combined with existing 10x scRNA-seq and 10x snATAC-seq, and performed integrative analysis of gene expression and DNA accessibility in human LAM lungs. We annotated cell types by mapping cells to the newly released LungMAP CellRef reference^19^. This revealed 33 different cell types present in the LAM lung, including the previously identified LAM^CORE^ cell population (**Fig. 1A & 1B**). Heatmaps showed highly concordant results for paired LAM snATAC-seq and scRNA-seq data (**Fig. 1B**). The combination of single cell ATAC and RNA data allowed us to identify cell-specific chromotin accessibility peaks and enriched transcription factor (TF) binding sites or motifs in a cell type specific manner. We found that LAM^CORE^ cell specific ATAC-seq peaks accessible regions were highly enriched motifs for the uterine selective homeobox family of transcription factors including PBX1, PBX3, HOXA9, HOXA10 and HOXD11. In addition, MITF, STAT family (STAT1, and 3), NR2F1, NR2F2, and TEAD family (TEAD 1, 2 and 3) were also listed as top enriched motifs in LAM^CORE^ cell accessible regions. Representative top enriched motifs and the corresponding TF expression in LAM vs. Control were shown in **Fig. 1C & 1D**. Among these, HOXD11, MITF and TEAD2 were selectively induced in LAM^CORE^ cells (pink bar) while PBX1, STAT3, and STAT3 were expressed in multiple cell types including LAM^CORE^ cells and their average expression levels are increased in LAM vs control (blue bar). MITF is a known driver oncogene for kidney angiomyolipoma and NR2F2 is a nuclear receptor that was recently identified by GWAS as a modifying genetic locus in LAM ^20^. The LAM^CORE^ cell specific peak nearby gene analysis identified uterine TFs including PBX1, PITX2, ESR1 and HAND2; many genes encoding muscle and colleagen (ACTA2, MYH11, COL1A2, and COL4A1) and genes involved in apoptosis selectively expressed in the LAM^CORE^ cell peak regions with increased expression in LAM vs. Control (**Fig. 1D & 1E**). We analyzed peak-to-gene (P2G) links in genomic regions of *HOXA10* and *PBX1* (**Fig. 1F**). As shown, there are uniquely accessible peaks detected in the promoter regions of *HOXA10* and 11 in LAM^CORE^ cells, their gene expression is uniquely induced in LAM^CORE^ cells as well. PBX1 has accessible peaks and induced expression in LAM^CORE^ cells, but neither the peak nor the gene expression is unique to LAM^CORE^ cells. As a cofactor of HOX family of TFs, the PBX1 selectivity to LAM^CORE^ cells is from the dimerization with HOX proteins. Taken together, the integrative scRNA/snATAC data analyses indicated the activation of uterine-specific transcriptional programming (i.e., homeobox family of transcription factors) at both gene expression and epigenetic level, and strongly support the notion that HOX-PBX signaling plays an important role in the transcription and epigenetic regulation of LAM lung pathogenesis.

**Figure 1.**
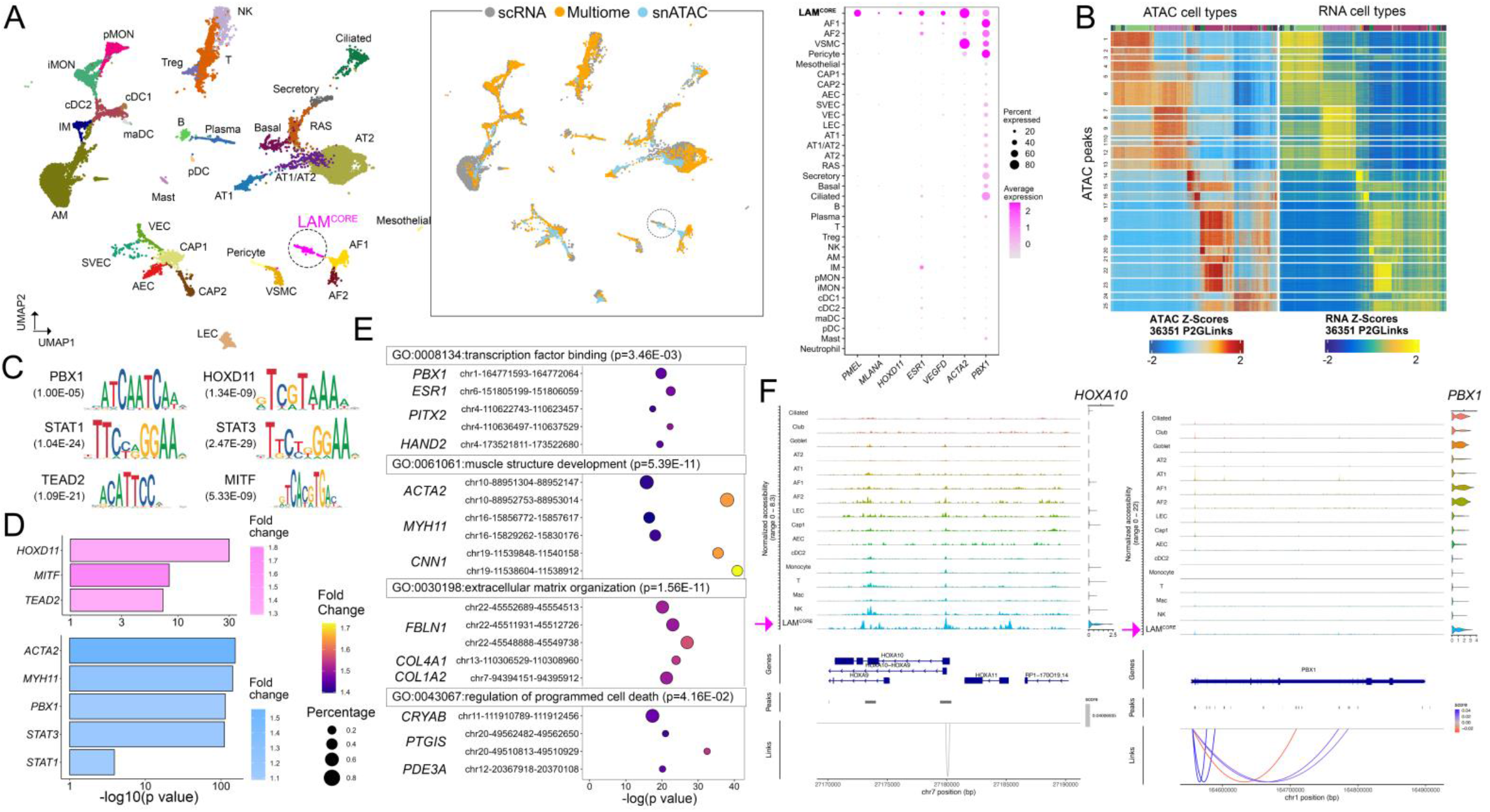
Integrative analysis of Single-cell transcriptome and chromatin accessibility in LAM lungs. (**A**) Left: UMAP visualization of integrated transcriptomic and epigenetic profiling of LAM lungs. The integration included 12,374 cells from scRNA-seq profiling and 10,554 nuclei from multiome and snATAC profiling. Cells/nuclei were colored by predicted cell types. Middle: UMAP visualization of cells/nuclei colored by data modalities and profiling methods. Right: Dotplot visualization of expression of representative genes differentially expressed in LAM^CORE^, including known LAM markers (PMEL, MLANA, VEGFD, ACTA2) and our predicted signature (HOXD11, ESR1, PBX1). (**B**) Heatmap visualization of identified peak-to-gene links (y axis) which had correlated ATAC accessibility and RNA expression patterns across cells (x axis) in RNA and ATAC profiling of LAM. Labels on the right represent k-means clusters of peak-to-gene links. (**C**) Motifs enriched in the differentially accessible peaks (DAPs) in LAM^CORE^ cells. (**D**) The expression levels of corresponding transcription factors and genes are increased in scRNA-seq of LAM vs. normal lung. Pink: Genes selectively induced in LAM^CORE^ ; Blue: Genes induced in LAM but not uniquely expressed in LAM^CORE^. (**E**) Gene Ontology terms enriched by genes near (<10kb) differentially accessible peaks (DAPs) in LAM^CORE^ cells. (**F**) Peak-to-gene (P2G) links in genomic regions of *HOXA10* (left) and *PBX1* (right). Tracks on the top left of each panel showed normalized ATAC accessibility in each cell type. Violin plots on the top right of each panel showed expression patterns of HOXA10 and PBX1 in each cell type. Scores of P2G links represent correlations of ATAC peak accessibility and RNA expression across cell types.

### Integrative analyses of PBX1 ChIP-seq and scRNA-seq of LAM lung identifies HOXs

PBX1 was reported as a pioneer factor that opens chromatin to recruit estrogen receptor alpha (ERα) in breast cancer cells ^21^. PBX (1-4) and HOX (1-11) are homeodomain TFs that form functional dimers and play important roles in different types of cancers ^12^. We applied integrative analyses of breast cancer PBX1 ChIP-seq (GSE28008) and scRNA-seq of LAM lung ^9^ to identify the potential PBX1 targets in LAM^CORE^ cells. Genes with positive PBX1 binding peaks and selectively expressed in LAM^CORE^ cells were identified, including *HOXD11, PITX2, ESR1, and NR2F2* (**Fig. 2A**). *STAT3* is predicted as a PBX1 transcriptional target with ubiquitous expression patterns in LAM and control lung cells, and its overall expression level is significantly higher in LAM vs. control lung (Figure 1E). Genes with positive PBX1 binding peaks that are also selectively expressed in LAM^CORE^ cells were functionally enriched in pathways relevant to LAM, including estrogen, ERK/MAPK and mTOR signaling (**Fig. 2B**), suggesting that PBX1 is directly involved in the regulation of genes and pathways leading to LAM pathogenesis.

**Figure 2.**
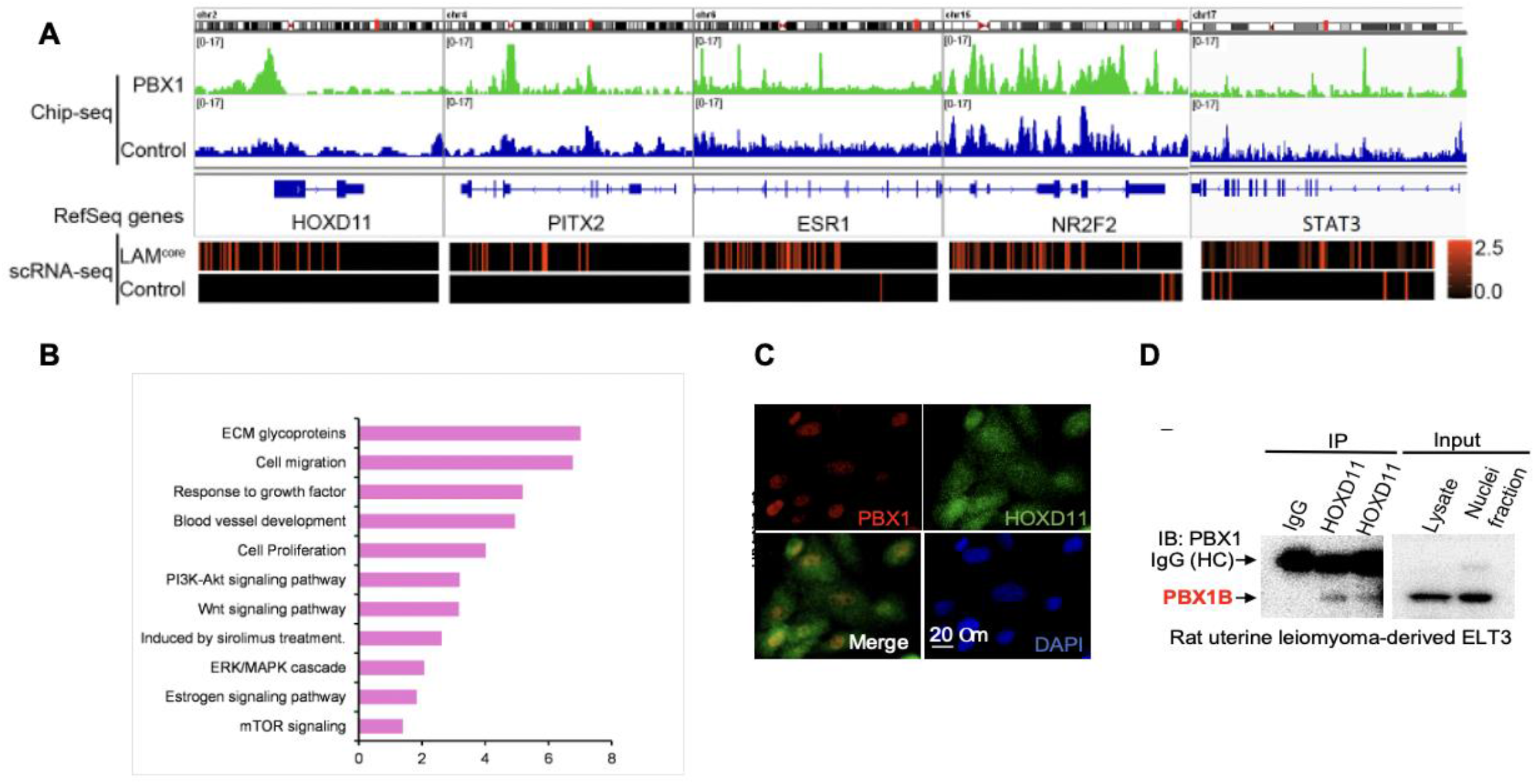
Integrative analyses of breast cancer PBX1 ChIP-seq and scRNA-seq of LAM lung identified PBX1-target genes in LAM lung cells. PBX1 interacts with HOXD11 in TSC2-null cells. (**A**) PBX1 targets selectively expressed in LAM^CORE^ cells were identified. (**B**) Functional enrichment of PBX1 targets in LAM^CORE^ cells. (**C**) Immunofluorescent staining of PBX1 and HOXD11 were performed in 621-101 cells. Confocal microscopy shows nuclear co-staining of PBX1 and HOXD11. (**D**) Nuclear fraction was isolated from ELT3 cells. Immunoprecipitation (IP) was performed using two HOXD11 antibodies (Sigma and CST). Mouse IgG was used as a negative control. Protein levels of PBX1 was assessed by immunoblotting in HOXD11-IP complex, protein lysate, and nuclear fraction of ELT3 cells.

### PBX1 interacts with HOXD11 in TSC2-null cells

Different members of PBX/HOX dimers determine tissue and target specificity. What kind of PBX/HOX dimer combinations are present in LAM lung is unknown. Our integrative scRNA-seq and snATAC-seq analysis identified highly selective expression of and chromatin accessibility for *PBX1, HOXA11* and *HOXD11* in LAM^CORE^. To determine the presence of specific HOX/PBX1 interaction, we performed immunofluorescence co-staining in LAM-derived 621-101 cells. Confocal microscopy showed that PBX1 immunoreactivity is concentrated in the nuclear compartment of TSC2-null cells (**Fig. 2C**). HOXD11 immunoreactivity is also prominent in the nucleus. Importantly, co-localization of PBX1 and HOXD11 is evident in the nucleus of TSC2-null LAM patient-derived 621-101 cells. To further demonstrate the interaction of PBX1 and HOXD11, we performed immunoprecipitation using two independent anti-HOXD11 antibodies, and detected PBX1 in the HOXD11 immunoprecipitation complex, protein extracts and nuclear fractions (**Fig. 2D**). Collectively, our findings indicate the interaction of PBX1 and HOXD11 in TSC2-null cells.

### Determine genomic and epigenetic regulation of LAM^CORE^ cells via transcriptional regulatory networks

To reveal the genomic and epigenomic regulatory circuits, predict crucial factors that determine LAM cell-specific cell fate, we constructed a LAM^CORE^ cell transcriptional regulatory networks (TRN) using PECA, a statistical model inferring context-specific TRN from paired gene expression and chromatin accessibility data. The model calculates the probability of the distribution of the expression of target genes conditional on the accessibility of regulatory elements and the expression of chromatin regulators, taking into consideration the activity status of regulatory elements and the expression of target gene. To ensure that the TRN is LAM^CORE^ specific, we require that TFs are abundantly expressed in LAM^CORE^ cells, and that target genes are LAM^CORE^ signature genes and genes induced in LAM^CORE^, compared to control mesenchymal cells. **Figure 3A** shows TFs with the most interactions in the LAM^CORE^ TRN (top 1% of the network). Hormone receptors *PGR* (progesterone receptor) and *AR* (androgen receptor), and homeobox TFs HOXA10, HOXA11, HOXD11, PBX1 and PBX3 are among the top ranked TFs in LAM network. Since PBX1 is a cofactor of HOX, forming functional dimers with HOX proteins ^12^, and we have demonstrated that PBX1 binds to HOXD11 in human and rat TSC2-null cells (Fig. 2C and 2D), we next focused on a PBX1-HOXD11 centered TRN by extracting out gene nodes directly connect to both PBX1 and HOXD11 in the network which include common upstream regulators, co-factors, and targets (**Fig. 3B**). PBX1-HOXD11 common targets were functionally enriched in “muscle proliferation/development”, “cell fate commitment”, “wound healing”, “response to hormone”, “Wnt signaling pathway”, “apoptotic signaling pathway” and “focal adhesion: PI3K-Akt-mTOR-signaling pathway” (**Fig. 3C)**. We also identified PBX-STAT common targets, and PBX1 and 3 edges were combined as PBX family and STAT 1 and 3 were combined as STAT family (F**ig. 3D**). Enriched bioprocess and signaling pathways of the PBX-STAT common target genes are shown in **Fig. 3E**.

**Figure 3.**
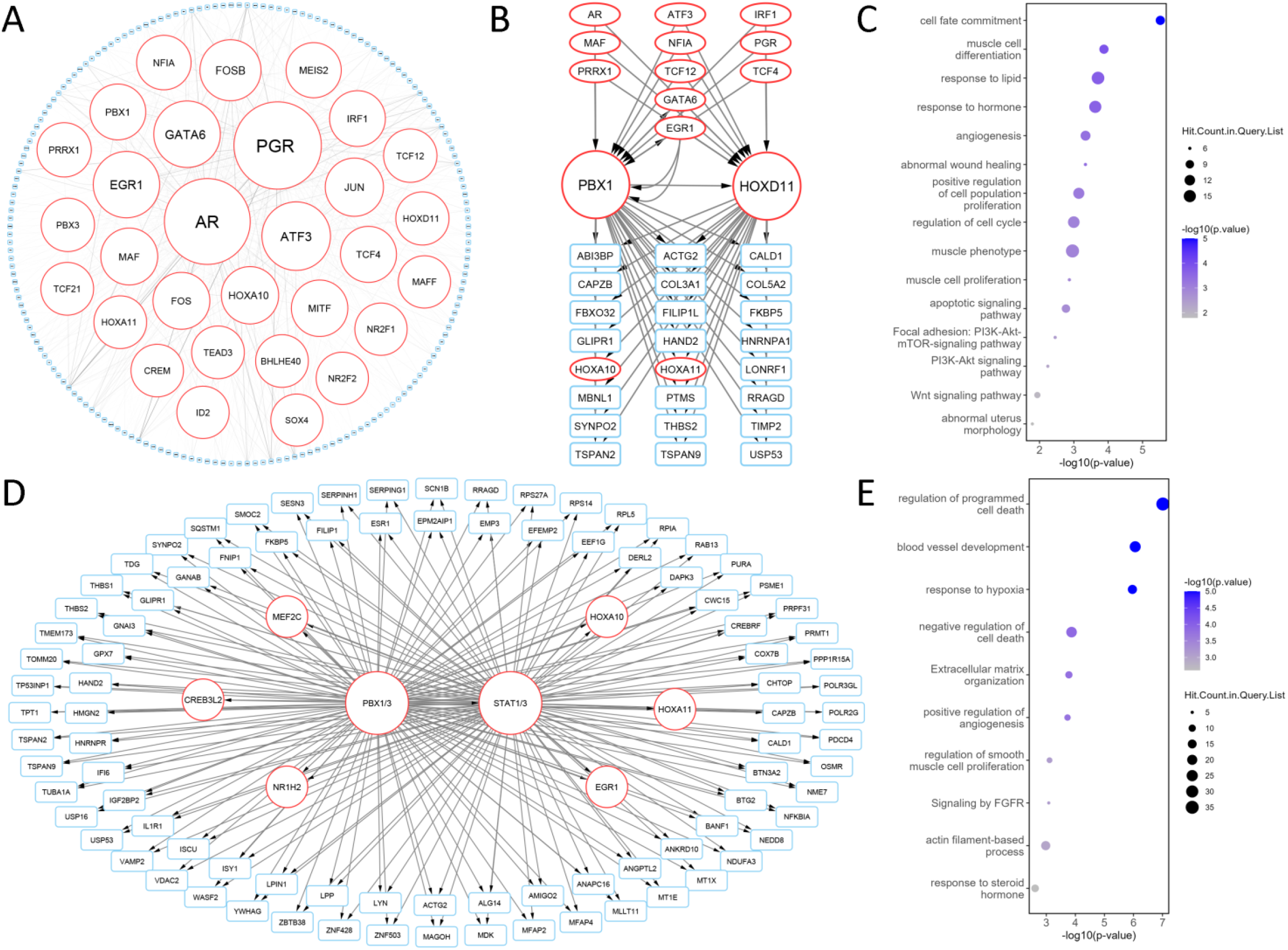
Transcriptional regulatory networks analysis using paired gene expression and chromatin accessibility data. **(A**) LAM^CORE^ cells transcriptional regulatory network (TRN) was constructed using PECA (top 1% of the network was displayed). LAM^CORE^ cells scRNA-seq of 4 LAM patients and snATAC-seq data from 3 LAM patients were used for the network construction. The edge regulation score cutoff was 99^th^ percentile. TF node size and font size are proportional to the number of edges connected to the given TF. Networks and connections were visualized using the Cytoscape (version 3.7.2) software package. (**B**) Identification of HOXD11-PBX1 common upstream regulators and downstream targets. (**C**) Enriched bioprocess and signaling pathways of the HOXD11-PBX1 network target genes. (**D**) Identification of PBX-STAT common targets. PBX1 and 3 edges were combined as PBX family and STAT 1 and 3 were combined as STAT family. (**E**) Enriched bioprocess and signaling pathways of the PBX-STAT common target genes.

### Expression of PBX1 and target genes is prominent in LAM

We first validated the expression of *PBX1* in LAM lung tissue (n=5) vs. non-LAM female lung (n=3). RT-PCR analysis showed a six-fold increase of *PBX1* transcript levels in LAM vs. Non-LAM-Female donor lung samples (p < 0.05) (**Fig. 4A**). Moreover, we re-analyzed a publicly available gene expression array dataset of laser-capture microdissected LAM lung cells and observed a significant increase of *PBX1* transcript levels in LAM (n=14) vs. female lung cancer samples (n=38) (p < 0.0001) (**Fig. 4B**). Immunohistochemical staining showed that PBX1 localizes to the nuclei of smooth-muscle actin (ACTA2)-positive LAM lung nodules from three LAM subjects (**Fig. 4C**). Furthermore, we observed positive PBX1 immunohistochemical staining in LAM uterine lesion cells (**Fig. 4D**) and renal angiomyolipoma cells (**Fig. 4E**). Importantly, in LAM lung tissue we found increased positivity and nuclear localization of the PBX1 transcriptional targets phospho-STAT1 and phospho-STAT3 (**Fig. 4F**). Together, our data show elevated expression of PBX1 at both transcript and protein levels, and activation of STAT1/3, consistent with the single cell multiomics findings.

**Fig. 4.**
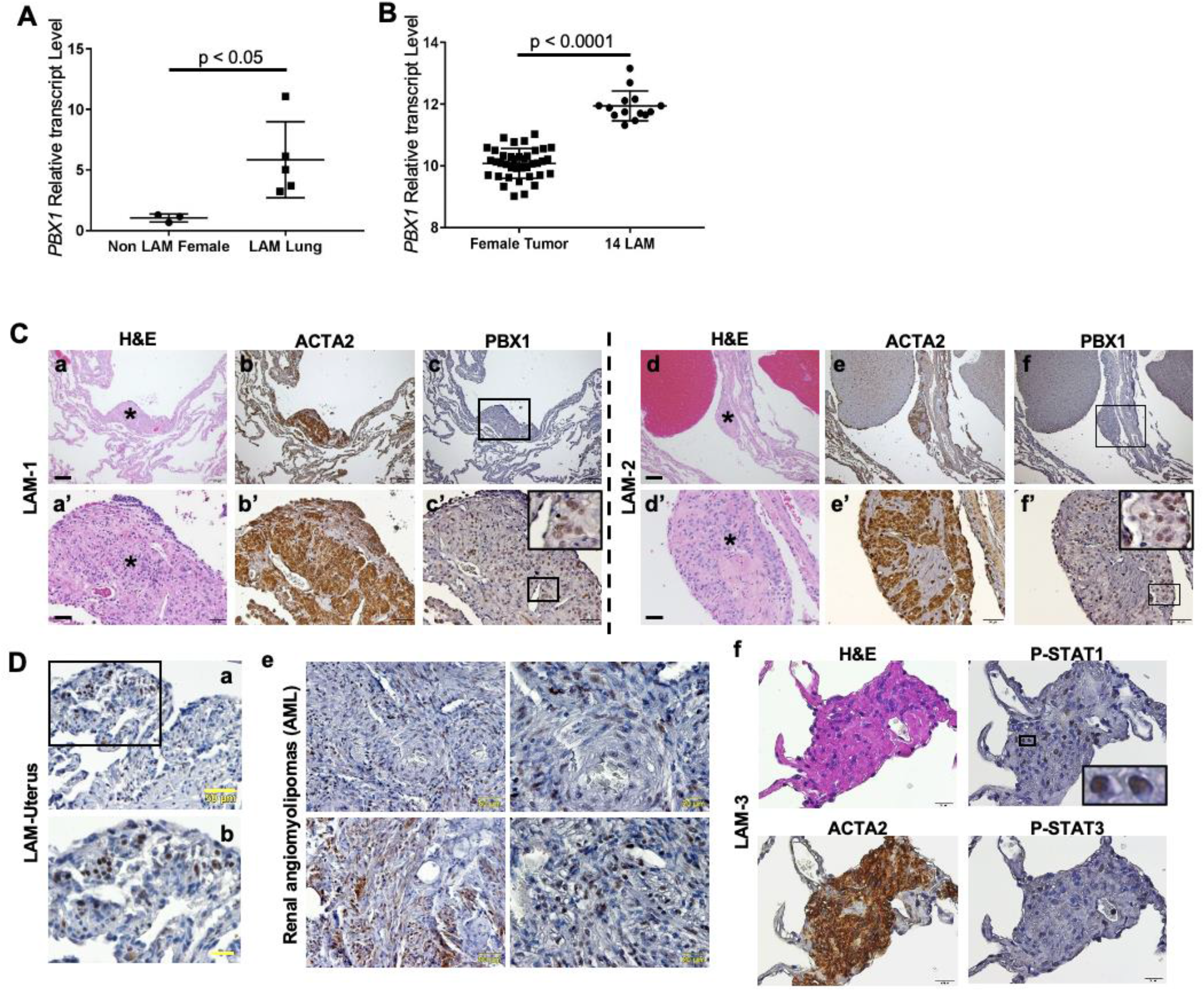
Expression of PBX1 is prominent in LAM lungs. (**A**) RT-qPCR for *PBX1* transcript levels in non-LAM female lungs (n=3) and LAM lungs (n=5). (**B**) Re-analysis of gene expression arrays of LAM lungs (n=14) and female tumors (n=38) shows the relative levels of *PBX1* transcript. p<0.05, p<0.0001, Student t-test. (**C**) Lung FFPE specimens from two LAM patients (LAM-1, a-c’; LAM-2, d-f’) were stained with H&E (a, a’, d, d’), smooth muscle actin (ACTA2) (b, b’, e, e’), and PBX1 (c, c’, f, f’). Scale bar 200 μm for a-f, and 50 μm for a’-f’. Asterisks in a, a’, d, and d’ indicate LAM nodules. (**D**) PBX1 Immunohistochemical staining of FFPE specimen from uterine tumor of a LAM patient. Scale bar 50 μm for a, 20 μm for b. (**E**) PBX1 immunohistochemical staining of FFPE renal angiomyolipoma specimens. (**F**) H&E and immunohistochemical staining for smooth muscle actin (ACTA2), phospho-STAT1 (Ser727), and phospho-STAT3 (Ser727) in FFPE specimen from a LAM lung.

### TSC2 negatively regulates PBX1 expression and STAT1/3 activation

To determine the molecular mechanisms underlying the upregulation of PBX1 expression and STAT1 activation, we first examined the effect of TSC2, a tumor suppressor gene that is frequently mutated and inactivated in LAM cells. We found that PBX1 protein levels were markedly increased by 7.8-fold in LAM patient-derived TSC2-null (TSC2-) relative to TSC2-addback cells (TSC2+) (**Fig. 5A and 5B**). We also found concomitantly higher levels of phospho-STAT1 (Ser727) by 3.1-fold and phospho-STAT3 (Ser727) by 5.8-fold in TSC2-null cells relative to that in TSC2-addback cells (**Fig. 5A and 5B**). Next, we examined PBX1 protein levels in xenograft tumors of Tsc2-null ELT3-V3 and TSC2-reexpressing ELT3-T3 cells and found a marked increase of PBX1 accumulation in Tsc2-null ELT3-V3 tumor cells relative to ELT3-T3 cells (**Fig. 5C**). Importantly, in a lung metastatic mouse model that we previously developed ^22^ PBX1 accumulation was more prominent in estrogen-induced lung metastatic lesions, compared to placebo-treatment, but not in adjacent normal lung cells (**Fig. 5D**). Collectively, our data suggest an important role of TSC2 in suppressing PBX1 expression and STAT1/3 phosphorylation in LAM-derived cells both in vitro and in vivo.

**Figure 5.**
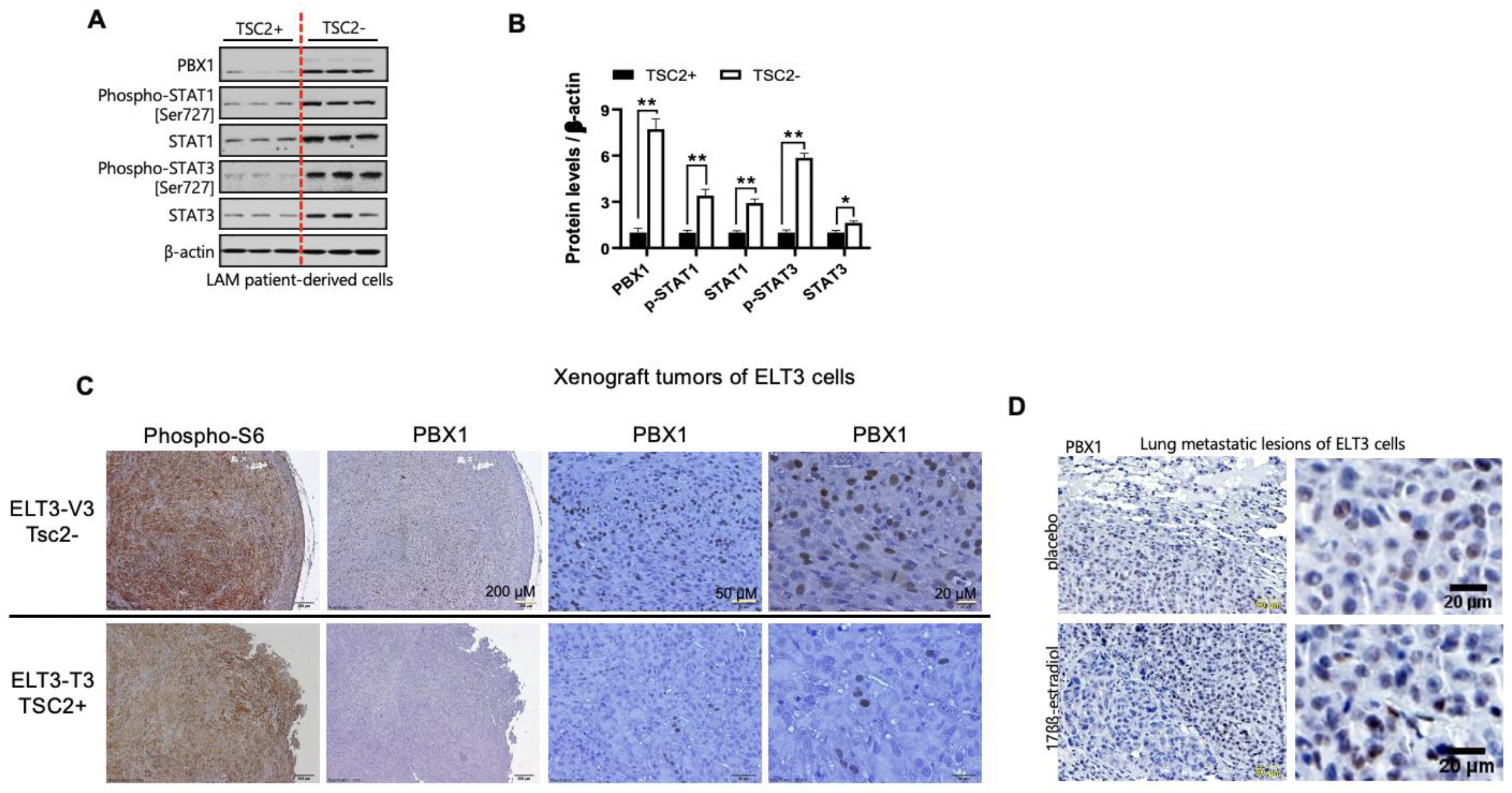
TSC2 negatively regulates the expression of PBX1 and the activation of STAT1/3. (**A**) Protein lysates were isolated from TSC2-null LAM-derived (TSC2-) and TSC2-addback cells (TSC2+) cells (n=3). Protein levels of PBX1, phosphp-STAT1 (Ser727), STAT1, phospho-STAT3 (Ser727), and STAT3 were assessed by immunoblotting analysis. β-actin as a loading control. (**B**) Densitometry analysis of protein levels normalized to β-actin. (**C**) 2×10^6^ Tsc2-null ELT3-V3 or TSC2-addback ELT3-T3 cells were subcutaneously injected into both flanks of 6-8 weeks female CB17Scid mice. Representative images of immunohistochemical staining of PBX1 in xenograft tumors. (**D**) 2×10^6^ Tsc2-null ELT3 cells were subcutaneously injected into both flanks of 6-8 weeks ovariectomized female CB17/Scid mice supplemented with placebo or 17β-estradiol. Representative images of immunohistochemical staining of PBX1 in xenograft tumors and lung metastatic lesions of ELT3 cells.

### Suppression of PBX1 attenuates STAT1/3 phosphorylation in LAM-derived cells

Since STAT family TFs (STAT1 and STAT3) is predicted as the most enriched motifs in the cell type specific chromatin accessibility regions of LAM^CORE^ cells (Fig. 1C) and as a direct transcription target of PBX1 (Fig. 2A), we further constructed PBX1/3 and STAT1/3 common network to identify common targets (**Fig. 6A**). Since families of TFs largely share TF binding sites or motifs, we combined PBX1 and 3 as PBX family and STAT1 and STAT3 as STAT family. Interestingly, “regulation of programmed cell death” is the most enriched function for PBX/STAT common target genes, and most of these genes are associated with “negative regulation of cell death” (**Fig. 6B**). Our network analysis suggested that PBX and STAT family of TFs work coherently to suppress LAM cell apoptosis.

**Figure 6.**
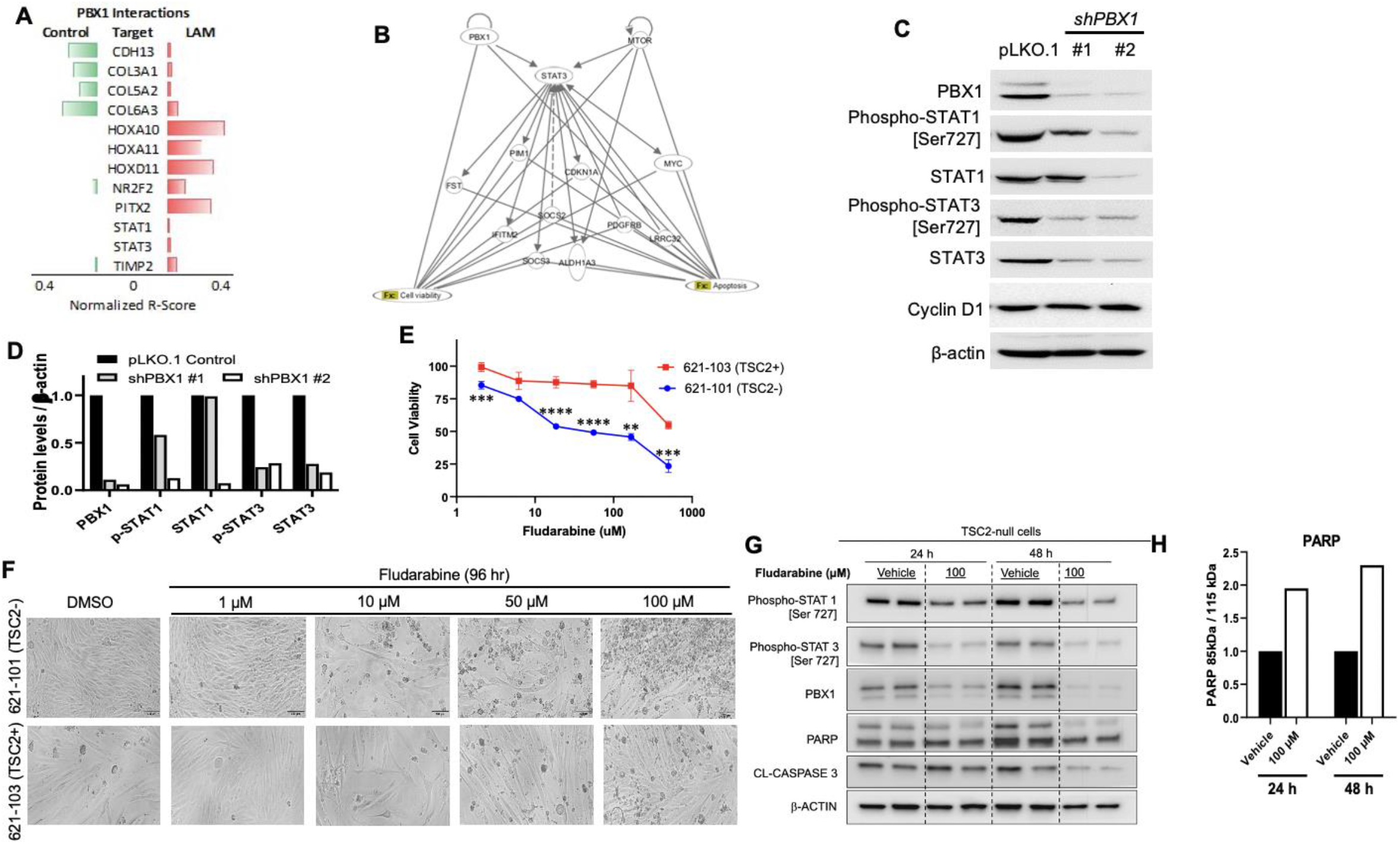
Suppression of PBX1 attenuates STAT1/3 phosphorylation and induces the death of TSC2-null LAM-derived cells. (**A**) Construction of PBX1/3 and STAT1/3 common network to identify common targets. (**B**) We combined PBX1 and 3 as PBX family and STAT 1 and 3 as STAT family and enriched function assessment of TF targets show negative regulation of cell death”. (**C**) TSC2-null LAM patient-derived 621-101 cells were infected with lentivirus of two independent PBX1-shRNAs or control pLKO.1 empty vector. Immunoblotting analysis of PBX1, phospho-STAT1 (Serine 727), STAT1, phospho-STAT3 (Serine 727), STAT3, and cyclin D1. β-actin as a loading control. (**D**) Densitometry analysis of protein levels normalized to β-actin. (**E**) Viability of TSC2-null 621-101 and TSC2-addback 621-103 cells (n=6 wells/treatment group) was assessed 72 hours after treatment with escalating concentrations of fludarabine. GI^50^ was calculated using CalcuSyn Software. **, ***, **** indicate statistical significance at P < 0.01, P < 0.001 and P < 0.0001, respectively (two-tailed unpaired Student’s T-test) (**F**) Phase-contrast microscopy was done to assess cell morphology after fludarabine treatment in 621-101 cells relative to 621-103 cells. (**G**) Immunoblotting analysis of TSC2-null 621-101 cells treated with fludarabine for 24 or 48 hours. Protein levels of phospho-STAT1(Ser727), phospho-STAT3 (Ser727), PBX1, and the cell death marker PARP cleavage was assessed. β-actin was used as a loading control.

To assess the impact of PBX1 on its downstream targets, we depleted PBX1 from 621-101 cells using lentiviral transduction with two different PBX1 shRNAs to avoid off-target effects and used pLKO.1 empty vector as a non-targeting control. Gene silencing was assessed by PBX1 immunoblotting. Densitometry assessment showed 90% and 95% reduction of PBX1 protein levels, compared to pLKO.1 control (**Fig. 6C and 6D**). Concomitantly, PBX1 silencing resulted in attenuation of phospho-STAT1 (Serine 727) by 45% and 88%, and phospho-STAT3 (Serine 727) by 80% and 75%, relative to pLKO.1 control. However, the protein level of the cell cycle protein Cyclin D1 was not affected by PBX1 depletion. Together, our results suggest that PBX1 activates STAT1 and STAT3 in Tsc2-null LAM patient-derived cells.

### Inhibition of STAT1 promotes the death of TSC2-null LAM-derived cells in vitro

To test if TSC2-null cells are sensitive to pharmacological suppression of STAT1, we treated LAM patient-derived cells with a STAT1-specific inhibitor fludarabine (NSC 118218, FaraA, Fludarabinum) that causes a specific depletion of STAT1 mRNA and protein (Nature Medicine volume 5, pages444– 447 (1999)). Fludarabine significantly decreased the viability of TSC2-null 621-101 cells (GI_50_=56.1 μM), compared to TSC2-expressing 621-103 cells (GI_50_=852.2 μM) within 72 hours of treatment **(Fig. 6E)**. In a separate experiment, we treated both TSC2-null 621-101 and TSC2-addback 621-103 cells (TSC2+) with escalating concentrations of fludarabine for 96 hours. Phase-contrast microscopy showed that 50 µM and 100 µM fludarabine treatment caused more profound cell death in 621-101 cells relative to that in 621-103 cells (**Fig. 6F**). Furthermore, fludarabine treatment for 48 hours resulted in marked reduction of phospho-STAT1 levels relative to vehicle control (**Fig. 6G**). Interestingly, fludarabine treatment reduced the protein levels of PBX1. Importantly, fludarabine treatment induced marked PARP cleavage in TSC2-null 621-101 cells, compared to vehicle treatment **(Fig. 6H)**, indicative of apoptosis induction by STAT1 inhibition. Collectively, our network analysis and functional studies consistently demonstrate that PBX1 and STAT family of TFs work coherently to suppress LAM cell apoptosis.

### Pharmacologic and molecular suppression of PBX1 promote the death of TSC2-null cells *in vitro* and *in vivo*

To assess the effect of PBX1 suppression on cell survival, TSC2-null 621-101 and ERL4 cells were treated with increasing concentrations of the PBX1 antagonist HXR9, a cell permeable synthetic peptide which inhibits HOX-PBX interaction and promotes cell killing ^23^. Cell morphology was evaluated 2 or 3 hours post treatment. We found that 25 µM HXR9 caused massive death of 621-101 (**Fig. 7A**) and ELT3 cells (**Fig. 7B**), respectively, indicative of *in vitro* cytotoxic effect of PBX1 suppression. Next, we examined the effect of HXR9 on lung colonization of ERL4 cells. Mice pre-treated with HXR9 exhibited lower levels of lung colonization relative to control mice 24- and 48-hours post treatment **(Fig. 7C and 7D)**. To assess the possible benefit of molecular suppression of *PBX1* on lung colonization, we intravenously injected female SCID mice with LAM-derived 621L9 cells infected with two independent PBX1-shRNAs or control pLKO.1. The photon flux of the bioluminescent signal was quantified. Bioluminescent intensities at the chest regions were similar in mice inoculated with pLKO.1 or PBX1-shRNA cells within one hour (baseline). Importantly, bioluminescent intensities were significantly reduced in mice inoculated with PBX1-shRNA cells relative to pLKO.1 control cell at 24-hours and 48-hours post cell inoculation (**Fig. 7E and 7F**). Collectively, these data suggest that PBX1 regulates the survival and lung colonization of TSC2-null LAM patient-derived cells *in vivo*.

**Figure 7.**
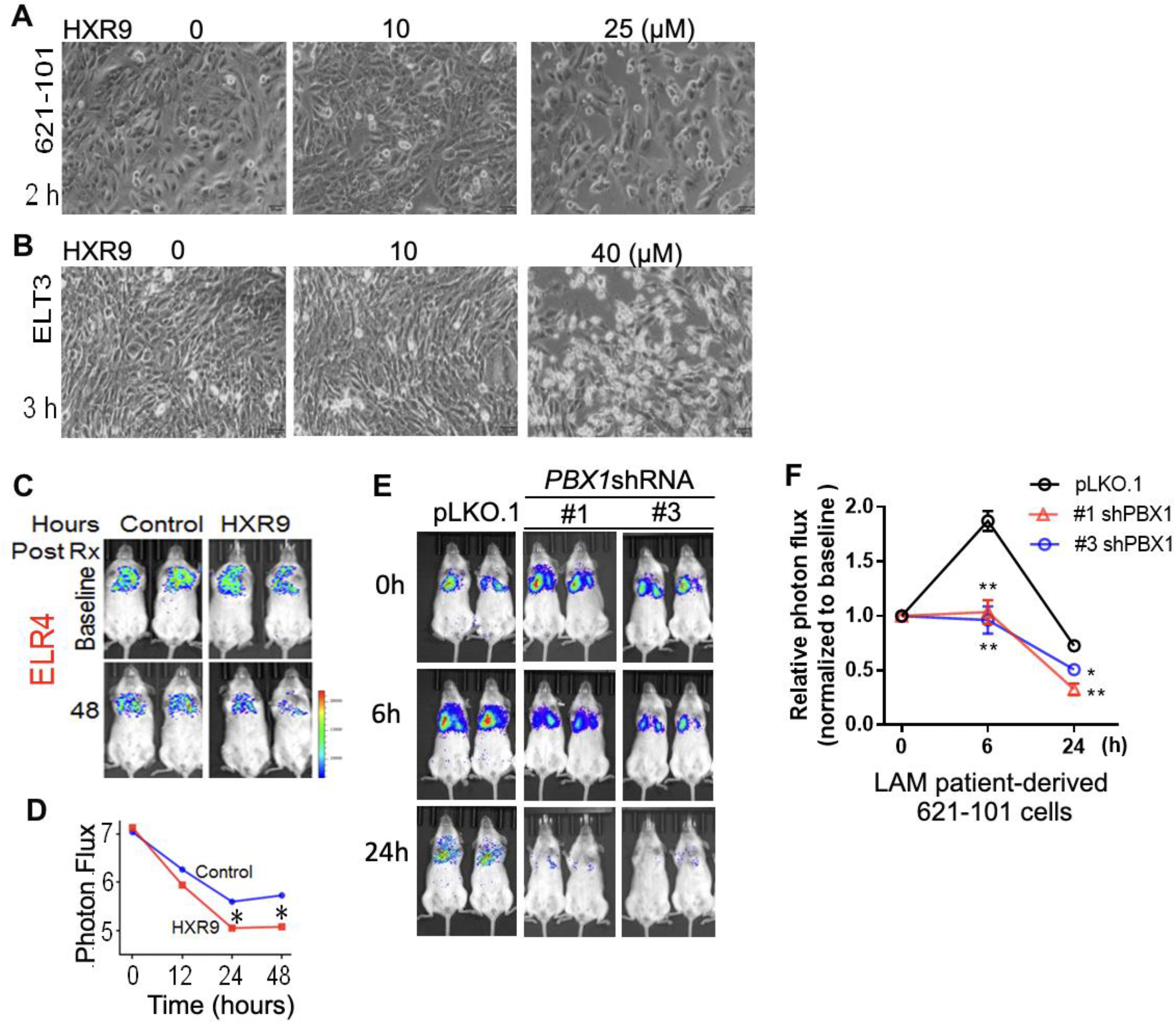
Suppression of PBX1 delays lung colonization of TSC2-null cells. (**A**) Tsc2-null rat uterine leiomyoma-derived ELT3 and (**B**) LAM patient-derived TSC2-null 621-101 cells were treated with PBX1 antagonist HXR9 with indicated concentration. Phase contrast light microscopy shows cell viability. (**C**) Female NSG mice were pre-treated with HXR9 (2 mg/kg, iv) for two days, and then intravenously inoculated with 2×10^5^ ERL4 cells. Bioluminescent imaging was performed. (**D**) Photon flux at chest regions was quantified. (**E**) 621-101 cells expressing luciferase (621L9) cells were infected with lentivirus of PBX1-shRNA cells or control pLKO.1 empty vector. Female NSG mice were intravenously inoculated with 621L9-shRNA-PBX1 or pLKO.1 control cells. (**F**) Bioluminescent imaging was performed at 0, 6 and 24 hours post cell-inoculation (n=4/group). * p<0.05, Student t-test.

## DISCUSSION

LAM has the strongest gender predisposition of any extra-genital human disease. Estrogen receptor (ESR1) and progesterone receptor (PGR) play an important role in LAM development and progression ^8, 24-26^. Clinical observations that the menstrual cycle influences symptoms and pneumothoraces, and that decline of lung function slows after menopause or oophorectomy and accelerates with exogenous estrogen (E2) use and pregnancy are consistent with this notion ^27-32^. We and others have reported that E2 promotes the survival and pulmonary metastasis of Tsc2-deficient cells in mouse models^7, 33-36^ and stimulates phosphorylation of pro-survival kinases MAPK and AKT in ELT3 (Eker rat uterine leiomyoma-derived)^37^ and in LAM patient-derived cells ^38 35, 39^. Human studies are difficult to perform with rigor because of the rare nature of the disease. Published retrospective case series of anti-hormone interventions have yielded conflicting results; the only LAM randomized trial that involved the aromatase inhibitor, letrozole, in postmenopausal women was under enrolled and was not adequately powered to address questions of efficacy (NCT01353209) ^12^. Given our convincing new data suggesting the uterine origin of LAM cells coupled with the existing data on treatment of uterine cancers, we have an opportunity to identify novel molecular targets for preclinical assessment that would assist in designing human anti-hormone regimens.

We observed the expression of uterine-specific HOXs (*HOXA11, HOXD11, EMX2) and* estrogen and progesterone receptors *(ESR1* and *PGR)* in pulmonary LAM^CORE^ cells that are not or rarely detectable in normal lung. We reasoned that HOXs and their targets are among the most dysregulated genes in GRN. We further identified significantly over-representative LAM^CORE^ signature genes containing positive PBX1 ChIP-seq peaks, suggesting HOX/PBX1 as a master regulator controlling the states transition of LAM^CORE^ cells. We performed scATACseq on LAM-lung and control donor lungs to determine and compare the alteration of chromatin accessibility states, conducted integrative analyses of scRNA-seq and scATACseq to identify HOX/PBX1 targets in LAM^CORE^ cells, constructed a LAM^CORE^ cell-specific GRN, characterized the role of HOX/PBX1 and determined the essential nodes in the GRN, and validated predicted direct HOX/PBX1 target genes by RT-PCR and immunohistochemistry.

Advances in single cell assays of scRNA-seq and scATAC-seq enable interrogation of important genomic/epigenomic aspects simultaneously which were previously impossible to address due to technological limitations. In this study, we exploit innovative technology to test novel ideas of direct clinical relevance including the prediction of the HOX/PBX1 regulatory circuit and testing the functional impact of perturbation of HOX/PBX1 network in cell-based LAM models. We also assess the potential therapeutic effect of HXR9 in LAM-relevant models *in vitro* and *in vivo*. We provide the first detailed genomic/epigenomic blueprint of HOX/PBX1 in distinct pulmonary LAM cells with the ultimate goal of understanding how HOX transcription factors, co-factors and network genes coordinate to regulate pulmonary LAM pathogenesis and progression, and provide proof-of-principle for targeting HOX/PBX1-regulated steroid-mediated transactivation to induce remission in LAM cells without affecting normal cells.

Functional enrichment analysis of PBX1 targets in LAM^CORE^ cells predicted the activation of pro-survival mTOR and ERK/MAPK signaling. We showed that PBX1 regulates LAM^CORE^ cell behaviors and metastatic potentials of TSC2-deficient cells mimicking findings in human LAM. We have previously demonstrated that E2 induced lung metastasis of ERL4 cells in SCID mice ^36^. In this study, we demonstrated functional impact of PBX1 on estrogen transactivation in vitro and in vivo. Our findings indicate a novel function of PBX1 in regulating the survival of TSC2-null cells.

As previously described [32], increased STAT3 phosphorylation is also seen in TSC2-null xenografted tumors. Goncharova EA, Goncharov DA, Damera G, Tliba O, Amrani Y, Panettieri RA, et al. STAT3 Signal transducer and activator of transcription 3 is required for abnormal proliferation and survival of TSC2-deficient cells: relevance to pulmonary lymphangioleiomyomatosis. *Mol Pharmacol*. 2009;76: 766–777. In serial sections, anti–phospho-JAK2 antibodies (Figure 2A) and anti–phospho-STAT3 (Ser-727) antibodies (Figure 2B) reacted with cytoplasmic proteins of spindle-shaped LAM cells. Nuclear reactivity with anti–phospho-STAT3 (Ser-727) antibodies (Figure 2B) was also seen in cells of LAM lung lesions. LAM lung lesions also reacted with anti–phospho-p44/42 antibodies (Figure 2C). Am J Respir Crit Care Med. 2010 Aug 15; 182(4): 531–539. We found that phospho-STAT1 (Ser72) is primarily localized in the nucleus of ACTA2-positive LAM cells, but not in neighboring lung cells. Phospho-STAT3 (ser72) is less prominent in LAM cell nucleus, indicating that STAT1 activation is an important event in LAM progression. Importantly, inhibition of STAT1 phosphorylation led to the death of TSC2-null cells but not TSC2-reexpressing cells.

## Acknowledgments

We thank The LAM Foundation and the National Diseases Research Interchange for assistance with tissue collection.

## METHODS

### Cell culture and reagents

Eker rat uterine leiomyoma-derived (ELT3) cells ^40, 41^ were kindly provided by Dr. C Walker, Institute of Biosciences and Technology Texas A & M University, Houston, TX. *Tsc2*^+/-^ A/J mice at 9 months old were obtained as a generous gift from Dr. Steve Roberds, Tuberous Sclerosis Alliance, Washington D.C. ELT3 cells stably expressing luciferase (ERL4) ^22^ and LAM patient-associated angiomyolipoma-derived (621-101) cells were kindly provided by Dr. EP Henske, Brigham and Women’s Hospital-Harvard Medical School, Boston, MA ^22, 42^. Cells were cultured in DMEM/F12 supplemented with 10% FBS, and 1% penicillin-streptomycin-amphotericin B (PSA).

### Cell viability assay

Cell viability was estimated using the 3-(4,5-dimethylthiazol-2-yl)-2,5-diphenyl tetrazolium bromide (MTT) assay. TSC2-null 621-101 and TSC2-addback 621-103 cells were plated in 96 well plates (2×10^3^/well). Cells were incubated in 37°C CO_2_ incubator for 24 hours prior to treatment with escalating concentrations of fludarabine (0-500 µM; #118218 Selleck Chemicals LLC, TX, USA) for 72 hours. 25 µL of MTT solution (2.5 mg/ml growth media) was added to each well, followed by 4h incubation of plates at 37°C. Formation of formazan crystals were observed under the microscope and solubilized with the addition of 125 µL of 0.04N HCl in isopropanol, followed by 90 minutes incubation of plates at 37°C. Cell viability was determined by measuring absorbance at 560 nm with a reference of 650 nm. Growth inhibitory power was analyzed using CalcuSyn Software {Chou, 2005 #965}{Chou, 1984 #964}

### Expression array analysis

Re-analysis of previously published expression array data (GEO accession number GSE16944) ^43^ was performed using an online tool GEO2R. Transcript levels of *PBX1* were compared between TSC2-deficient (TSC2-) and TSC2-addback (TSC2+) cells, or rapamycin-treated and vehicle-treated TSC2-deficient (TSC2-) cells.

### Quantitative RT-PCR

RNA from cultured cells was isolated using RNeasy Mini Kit (Qiagen). Gene expression was quantified using One-Step qRT-PCR Kits (Invitrogen) in the Applied Biosystems Step One Plus Real-Time PCR System, and normalized to beta-actin (human) or alpha-tubulin (rat). Primers used are listed in supplementary table.

### Immunofluorescence staining

Sections were deparaffinized, incubated with primary antibody (1:100 in PBS+3%BSA) and smooth muscle actin (SMA, 1:200 in PBS+3% BSA, Santa Cruz, # sc32251), and secondary antibodies (1:1,000, Invitrogen, #A-21202 and #A10042). Images were captured with Fluorescence Microscope (Olympus BX60).

### Confocal Microscopy

ELT3 cells and LAM-derived cells were plated overnight on glass coverslips in 12-well tissue culture plates. Cells were serum starved overnight, and then treated with 20 nM rapamycin for 24 hours. Cells were rinsed with PBS twice, fixed with warm 4% paraformaldehyde, permeabilized with 0.2% Triton X-100, blocked in 3% BSA/PBS for 1 hr, and then incubated with primary antibody 1% BSA in PBS for 1 hour followed by secondary antibodies for 1 hour at room temperature. Images were captured with a FluoView FV-10i Olympus Laser Point Scanning Confocal Microscope.

### Phase-contrast microscopy

TSC2-null 621-101 and TSC2-addback 621-103 cells were seeded in 6-well plates (5×10^4^ cells/well) 24 hours prior to treatment with increasing concentrations of fludarabine (0-100 µM; Selleck Chemicals LLC, Tx, USA) for 96h. Cell morphology images were captured using Phase-contrast microscopy.

### Immunohistochemistry

Sections were deparaffinized, incubated with primary antibodies and biotinylated secondary antibodies and counterstained with Gill’s Hematoxylin.

### Immunoblotting

Protein lysis of TSC2-null 621-101 cells treated with fludarabine was done using mPER buffer (#78501 Thermo Fisher Scientific) containing protease and phosphate inhibitors. Lysates were resolved on 4–20% Mini-PROTEAN TGX Precast gels (Bio-Rad #4561094) and transferred using a wet electrophoretic transfer unit (Bio-Rad) onto PVDF membranes (Thermo Fisher Scientific). Membranes were incubated at 4°C overnight in primary antibody after 1h blocking with 5% non-fat dry milk. Primary antibodies include Phospho-STAT1 (#8826), Phospho-STAT3 (#9145), PBX1 (#4342), PARP (#9532), and β-Actin (#3700) from Cell Signaling Technology. Secondary incubation was done using Horseradish peroxidase-linked secondary antibody for 2 hours on a shaker. Signal was detected with SuperSignal West Pico PLUS Chemiluminescent Substrate (#PI34580 Thermo Fisher Scientific).

### shRNA downregulation

293T packaging cells were transfected with PBX1 shRNA, or non-Targeting shRNA vectors pLKO.1 using Mirus Trans-IT TKO Transfection reagent (Mirus). 621-101 cells were transduced with lentiviruses for 48 hours and then selected against puromycin. Stable clones were harvested for future experiments.

### Animal studies

Female C.B-*Igh-1*^*b*^/IcrTac-*Prkdc*^*scid*^ (SCID) (Taconic Biosciences, Germantown, NY) or NSG (Jackson Laboratory) mice at 6-8 week of age were used in this study. 2×10^5^ cells were injected into mice intravenously as previously described ^22^. Animal health was monitored daily during the tumor experiments. All mice were euthanized by carbon dioxide (CO_2_) inhalation via compressed gas after the last image was taken.

### Drug treatment *in vivo*

Mice inoculated with Tsc2-null cells were randomized and treated with the following two groups: CXR9 or HXR9 (2 mg/kg/day i.v.) for two days.

### Bioluminescent reporter imaging

Ten minutes prior to imaging, mice were given D-luciferin (120 mg/kg, i.p., PerkinElmer Inc., # 122799). Bioluminescent signals were recorded using the Xenogen IVIS Spectrum System. Total photon flux of chest regions was analyzed as previously described ^22^.

### Statistical analyses

Data represents mean ± SD. Statistical analyses were performed using two-tailed Student’s t-test when comparing two groups for in vitro and in vivo studies, and one way ANOVA test (Dunnett’s multiple comparisons test when comparing multiple groups with control group, Tukey’s multiple comparisons test when making multiple pair-wise comparisons between different groups) for multiple group comparisons. A *P* value less than 0.05 was considered significant.

### Study approval

The University of Cincinnati Standing Committees on Animals approved all procedures described according to standards as set forth in The Guide for the Care and Use of Laboratory Animals. The Institutional Review Board of the University of Cincinnati approved all human relevant studies.

## References

1. Astrinidis, A. et al. Mutational analysis of the tuberous sclerosis gene TSC2 in patients with pulmonary lymphangioleiomyomatosis. J Med Genet 37, 55–57 (2000).

2. Smolarek, T.A. et al. Evidence that lymphangiomyomatosis is caused by TSC2 mutations: chromosome 16p13 loss of heterozygosity in angiomyolipomas and lymph nodes from women with lymphangiomyomatosis. Am J Hum Genet 62, 810–815 (1998).

3. Carsillo, T., Astrinidis, A. & Henske, E.P. Mutations in the tuberous sclerosis complex gene TSC2 are a cause of sporadic pulmonary lymphangioleiomyomatosis. Proc Natl Acad Sci U S A 97, 6085–6090 (2000).

4. Bittmann, I., Rolf, B., Amann, G. & Lohrs, U. Recurrence of lymphangioleiomyomatosis after single lung transplantation: new insights into pathogenesis. Hum Pathol 34, 95–98 (2003).

5. Karbowniczek, M. et al. Recurrent lymphangiomyomatosis after transplantation: genetic analyses reveal a metastatic mechanism. Am J Respir Crit Care Med 167, 976–982 (2003).

6. Hayashi, T. et al. Prevalence of uterine and adnexal involvement in pulmonary lymphangioleiomyomatosis: a clinicopathologic study of 10 patients. Am J Surg Pathol 35, 1776–1785 (2011).

7. Prizant, H. et al. Uterine-specific loss of Tsc2 leads to myometrial tumors in both the uterus and lungs. Mol Endocrinol 27, 1403–1414 (2013).

8. Prizant, H. et al. Estrogen maintains myometrial tumors in a lymphangioleiomyomatosis model. Endocr Relat Cancer 23, 265–280 (2016).

9. Guo, M. et al. Single-Cell Transcriptomic Analysis Identifies a Unique Pulmonary Lymphangioleiomyomatosis Cell. Am J Respir Crit Care Med 202, 1373–1387 (2020).

10. Daftary, G.S. & Taylor, H.S. Endocrine regulation of HOX genes. Endocr Rev 27, 331–355 (2006).

11. Agarwal, S., Bell, C.M., Taylor, S.M. & Moran, R.G. p53 Deletion or Hotspot Mutations Enhance mTORC1 Activity by Altering Lysosomal Dynamics of TSC2 and Rheb. Mol Cancer Res 14, 66–77 (2016).

12. Alharbi, R.A. et al. Inhibition of HOX/PBX dimer formation leads to necroptosis in acute myeloid leukemia cells. Oncotarget 8, 89566–89579 (2017).

13. Carew, J.S., Kelly, K.R. & Nawrocki, S.T. Mechanisms of mTOR inhibitor resistance in cancer therapy. Target Oncol 6, 17–27 (2011).

14. Hewitt, S.C., Deroo, B.J. & Korach, K.S. Signal transduction. A new mediator for an old hormone? Science 307, 1572–1573 (2005).

15. Javed, S. & Langley, S.E. Importance of HOX genes in normal prostate gland formation, prostate cancer development and its early detection. BJU Int 113, 535–540 (2014).

16. Morgan, R. et al. Antagonism of HOX/PBX dimer formation blocks the in vivo proliferation of melanoma. Cancer Res 67, 5806–5813 (2007).

17. Pierard, G.E. & Pierard-Franchimont, C. HOX Gene Aberrant Expression in Skin Melanoma: A Review. J Skin Cancer 2012, 707260 (2012).

18. Guo, M. et al. Single Cell Transcriptomic Analysis Identifies a Unique Pulmonary Lymphangioleiomyomatosis Cell. Am J Respir Crit Care Med (2020).

19. Guo, M. et al. Guided construction of single cell reference for human and mouse lung. bioRxiv, 2022.2005.2018.491687 (2022).

20. Wirka, R.C. et al. Atheroprotective roles of smooth muscle cell phenotypic modulation and the TCF21 disease gene as revealed by single-cell analysis. Nature Medicine (2019).

21. Magnani, L., Ballantyne, E.B., Zhang, X. & Lupien, M. PBX1 genomic pioneer function drives ERalpha signaling underlying progression in breast cancer. PLoS Genet 7, e1002368 (2011).

22. Yu, J.J. et al. Estrogen promotes the survival and pulmonary metastasis of tuberin-null cells. Proc Natl Acad Sci U S A 106, 2635–2640 (2009).

23. Platais, C. et al. Targeting HOX-PBX interactions causes death in oral potentially malignant and squamous carcinoma cells but not normal oral keratinocytes. BMC Cancer 18, 723 (2018).

24. Prizant, H. & Hammes, S.R. Minireview: Lymphangioleiomyomatosis (LAM): The “Other” Steroid-Sensitive Cancer. Endocrinology 157, 3374–3383 (2016).

25. Chen, Q. et al. Xist repression shows time-dependent effects on the reprogramming of female somatic cells to induced pluripotent stem cells. Stem Cells 32, 2642–2656 (2014).

26. Logginidou, H., Ao, X., Russo, I. & Henske, E.P. Frequent estrogen and progesterone receptor immunoreactivity in renal angiomyolipomas from women with pulmonary lymphangioleiomyomatosis. Chest 117, 25–30 (2000).

27. Kinoshita, M. et al. Hormone receptors in pulmonary lymphangiomyomatosis. Kurume Med J 42, 141–144 (1995).

28. Ohori, N.P., Yousem, S.A., Sonmez-Alpan, E. & Colby, T.V. Estrogen and progesterone receptors in lymphangioleiomyomatosis, epithelioid hemangioendothelioma, and sclerosing hemangioma of the lung. Am J Clin Pathol 96, 529–535 (1991).

29. Johnson, S.R. & Tattersfield, A.E. Decline in lung function in lymphangioleiomyomatosis: relation to menopause and progesterone treatment. Am J Respir Crit Care Med 160, 628–633 (1999).

30. McCormack, F.X. Lymphangioleiomyomatosis: a clinical update. Chest 133, 507–516 (2008).

31. Taveira-DaSilva, A.M., Pacheco-Rodriguez, G. & Moss, J. The natural history of lymphangioleiomyomatosis: markers of severity, rate of progression and prognosis. Lymphat Res Biol 8, 9–19 (2010).

32. Taveira-DaSilva, A.M., Stylianou, M.P., Hedin, C.J., Hathaway, O. & Moss, J. Decline in lung function in patients with lymphangioleiomyomatosis treated with or without progesterone. Chest 126, 1867–1874 (2004).

33. Akbaraly, T.N. et al. Chronic inflammation as a determinant of future aging phenotypes. CMAJ 185, E763–770 (2013).

34. Liu, F. et al. Real-time monitoring of tumorigenesis, dissemination, & drug response in a preclinical model of lymphangioleiomyomatosis/tuberous sclerosis complex. PLoS One 7, e38589 (2012).

35. Huang, Y.S., Hsieh, H.Y., Shih, H.M., Sytwu, H.K. & Wu, C.C. Urinary Xist is a potential biomarker for membranous nephropathy. Biochem Biophys Res Commun 452, 415–421 (2014).

36. Diab, K.J. et al. Stimulation of sphingosine 1-phosphate signaling as an alveolar cell survival strategy in emphysema. Am J Respir Crit Care Med 181, 344–352 (2010).

37. Finlay, G.A. et al. Estrogen-induced smooth muscle cell growth is regulated by tuberin and associated with altered activation of platelet-derived growth factor receptor-beta and ERK-1/2. J Biol Chem 279, 23114–23122 (2004).

38. Jones, K.A., Jiang, X., Yamamoto, Y. & Yeung, R.S. Tuberin is a component of lipid rafts and mediates caveolin-1 localization: role of TSC2 in post-Golgi transport. Exp Cell Res 295, 512–524 (2004).

39. Araujo, E.S., Vasques, L.R., Stabellini, R., Krepischi, A.C. & Pereira, L.V. Stability of XIST repression in relation to genomic imprinting following global genome demethylation in a human cell line. Braz J Med Biol Res 47, 1029–1035 (2014).

40. Everitt, J.I., Wolf, D.C., Howe, S.R., Goldsworthy, T.L. & Walker, C. Rodent model of reproductive tract leiomyomata. Clinical and pathological features. Am J Pathol 146, 1556–1567 (1995).

41. Howe, S.R., Gottardis, M.M., Everitt, J.I. & Walker, C. Estrogen stimulation and tamoxifen inhibition of leiomyoma cell growth in vitro and in vivo. Endocrinology 136, 4996–5003 (1995).

42. Hong, F. et al. mTOR-raptor binds and activates SGK1 to regulate p27 phosphorylation. Mol Cell 30, 701–711 (2008).

43. Krishnakumar, R. & Kraus, W.L. The PARP side of the nucleus: molecular actions, physiological outcomes, and clinical targets. Mol Cell 39, 8–24 (2010).

